# Male *Anolis* lizards prefer virgin females

**DOI:** 10.1101/421925

**Authors:** Jessica Stapley

## Abstract

Males can gain fitness benefits by preferentially courting and mating with virgin females if they represent a lower risk of sperm competition. When females mate multiply, but do not remate frequently, males can experience a lower level of sperm competition when mating with virgins. Male preference for virgins has been demonstrated many times in invertebrates but rarely in vertebrates. In this study, I tested if *Anolis apletophallus* males preferentially courted virgin females more than non-virgin females, in two-choice trails where the virgin was smaller than the non-virgin, and trails where the females were size-matched. In both trails males preferentially courted the virgin females more than non-virgin females. This suggests males can discriminate between females based on their reproductive history and that they do not use body size as a cue. Males most likely used visual signals from the female, although these signals could not be identified in this study. This is only the second study to show male preference for virgins in vertebrates. Although it is possible that male preference for virgins is relatively rare in vertebrates, I argue that certain life history traits, namely, where large females do not have reproductive benefits, and or when sperm is stored between subsequent reproductive events this preference function can evolve. Future studies focusing on such systems are likely to be fruitful with respect to this male mating preference and may help us to better understand the evolution of male mating preferences and female traits.

## Introduction

Although, females are generally considered the choosier sex, male mating preferences have also been demonstrated in a range of taxa when male courtship is costly and when female quality varies (for review see Bonduriansky 2001). The most common male mate preference reported is preference for larger, more fecund females (insects (Bonduriansky 2001), lizards (Whiting & Bateman 1999), snakes (Shine et al. 2003); and fish (Ptacek & Travis 1997; Wong & Jennions 2003). Amongst invertebrates another common male mating preference is preference for virgins (Bonduriansky 2001). Male mate preference for virgins is likely to maximize a male’s fertilization success because of a reduction in sperm competition (see Bonduriansky 2001). Despite considerable evidence that males discriminate between females based on reproductive history in invertebrates, this has only been demonstrated once in a vertebrate, in the guppy (Guevara-Fiore et al. 2009).

The dearth of studies in vertebrates demonstrating male preference for virgins begs the question; is male preference for virgins less common in vertebrates or does this simply represent a taxonomic bias in previous studies? For males to gain benefits from reduced sperm competition by mating with females, certain conditions are needed and some of these may be less common in vertebrates. Firstly, it requires internal fertilization, a common feature of vertebrates, however this precludes many fish species and amphibians. The one vertebrate example of male mate preference for virgin females comes from the guppy, which is one of the few fish that has internal fertilization (Guevara-Fiore et al. 2009). A preference for virgins is unlikely to evolve when larger or older females represent greater reproductive value, and this is more likely in long-lived species. As already mentioned when larger females are more fecund males often prefer to mate with larger females (e.g. Bonduriansky 2001; Wong & Jennions 2003). A recent study in guppies found that male preference for larger females can persist in populations even when fecundity is not correlated to female body size, because populations had diverged in life history strategies in different environments (Arriaga & Schlupp 2013). When females continue to grow after maturation, a situation that is common in vertebrates but rare in invertebrates, then the fecundity benefits of larger, non-virgin females may outweigh any benefits of reduced sperm competition achieved when mating with smaller, virgin females. Likewise, if older females have higher reproductive success as is the case for many birds (e.g. Angelier et al. 2007) and mammals (Margulis et al. 2005), then a preference for less experienced females is unlikely to evolve. Males that preferentially mate with virgins will obtain few benefits if the female remates immediately after her first mating and or if female remating rate is high, because there will be only a short time window when sperm competition is reduced (Bonduriansky 2001). Finally, if reproduction spans multiple distinct breeding seasons, i.e. if species are iteroparus and females do not store sperm between these events, males should evolve strategies to choose females who have not mated in that breeding season rather than choosing individuals that have never mated. The evolution of male mating preference for virgins, thus may be dependent on life history - short longevity and low remating frequency will promote the evolution of preference for virgins.

Many of the conditions necessary for the evolution of male mating preference for virgins are present in the anolis lizard *Anolis apletophallus*. The species has a short life cycle going from egg to adult in ∼160 days, and with less than 5% annual survival in the field (Andrews & Nichols 1990), they are essentially an annual species. Females lay eggs every 7-10 days, which can be fertilized from stored sperm from previous matings (Andrews & Rand 1974). Clutches from field caught females had between 1-2 fathers (18 clutches, median clutch size =8) (J. Stapley unpublished data), so there is the potential for sperm competition. However, based on observations in the lab *A. apletophallus* seems reluctant to remate. For example, when placed with two males sequentially, each trial separated by 4 days, females never remated with the second male (J. Stapley unpublished data). Also, females that were not sperm depleted (producing fertile eggs in the lab) did not mate readily in mating trails (J. Stapley unpublished data). Thus, it appears that remating rates are not high in *A. apletophallus*, compared to other lizards (Calsbeek et al. 2007; Stapley 2008; Keogh et al. 2013). Finally, there is no evidence that female body size is related to fecundity. From females maintained in captivity (90 females, 956 eggs) female body size was not related to egg weight or hatchling weight, and inter egg interval was not related to female body size (J. Stapley unpublished data). In this species, clutch size is not a useful quantity to assess variation in fecundity, because clutch size is fixed at 2 in anoles, and in *A. apletophallus* the majority of eggs are laid as singletons. Thus, *A. apletophallus* is essentially an annual species that has a relatively low remating frequency. Males that preferentially mate with virgin females have 100% paternity assurance of the first eggs that the female produces, with the probability of paternity dropping fairly slowly. Mating with smaller, virgin females is unlikely to have cost in terms of reduced female fecundity because there is no relationship between female body size and fecundity.

Here I test if male *Anolis apletophallus* lizards prefer virgin females and if they use female body size to discriminate between virgins and non-virgins, as virgin females encountered by males in the field are smaller (<40mm) that non-virgin females(Andrews 1989). I conducted two experiments; in the first I tested if males discriminate between a small virgin and a large non-virgin (a naturally occurring situation) and if they preferred virgins. I predicted that males would preferentially court the virgin to maximize their fertilisation success. In the second experiment, I tested if males could discriminate between a pair of large size matched females when one was a virgin and the other was not. If males use body size as a cue to virginity I predicted that males would display equally to both females. The large virgins probably never occur naturally in the field, except in cases of very low density.

## Methods

Lizards were collected from Sobernia National Park and Barro Colarado Island in Panama. All lizards were housed in the laboratory for at least 4 weeks prior to experiments to allow them to acclimate. During this time they were housed individually in mesh cages (1m high X 60cm wide X 60xm long) that contained several dowel perches. These individual enclosures were housed within a large screened building on the edge of the rainforest. As such, light, temperature and humidity were the same as the surrounding rainforest. Animals were fed insects caught by sweep netting surrounding grass and rainforest edges, and lizards were sprayed with water daily. Enclosures were checked daily for eggs. Virgin females were small females (<40mm snout vent length (SVL)) when collected in the field and after being held individually in the laboratory produced an infertile egg – indicating sexual maturity. Larger virgin females (43-45 mm) were females that were caught before reaching sexual maturity and held in isolation in the laboratory. Non-virgins were larger females (>40mm) when caught in the field and who had produced fertile eggs in the laboratory. Egg fertility is easily distinguished based on egg colour and the presence of a leathery shell. Infertile eggs are yellow, soft and lack an obvious shell; fertile eggs are white, firm and have a calcified shell.

In the first experiment (November-January) 12 males chose between a small virgin female (mean SVL and standard error = 40 ± 0.21mm) and a larger non-virgin (mean SVL = 45 ± 0.38mm). In the second experiment (January-March) 14 males had a choice between size-matched (within 1mm) virgin and non-virgin females (mean SVL = 44 ± 023mm). In both experiments females and males were unfamiliar, they were collected from different populations approximately 1km apart and could not see each other while held in the laboratory. A total of 26 females were used, females were used twice but not in the same pair and had at least 10 days rest between experiments.

During the trials, males were placed into individual experimental cages (1m long X 1m high X 0.5m) where they remained for at least one week prior to experiments. This is long enough for lizards to acclimate and adopt normal behaviours such as movement between perches and assertion displays (J Stapley unpublished data). Each male enclosure contained two female enclosures at opposite ends of his enclosure, approximately 80 cm apart. The female enclosures were round (15cm diameter X 15 cm high) plexiglass enclosures with a single dowel perch. Male snout vent length ranged from 43-48mm (mean 44.71mm) in the first experiment and 42-47mm (mean 45.3mm) in the second trial.

On the day of the experiment each female was placed into one of the female enclosures and male behaviour was recorded for one hour with a digital video camera. During this time, the camera was trained on the male and as such female behaviour was not formally recorded or analysed. The placement of virgins and non-virgins was balanced so half of the virgins went in the left enclosure and half went into the right.

During analysis of the video, the number of times a male displayed to each female was recorded. The direction of the display is easily assigned because males display with their dewlap perpendicular to the receiver and often look at the receiver before and after the display. The number of displays was compared using a Wilcox Sign Rank test. As females were on opposite sides of the enclosure there was no ambiguity in identifying the direction of the display. The male’s position varied at the beginning of the experiment so I recorded if he was closer to the virgin or the non-virgin and tested if he preferentially displayed to the closer female using an Exact Binomial Test. The relationship between total number of displays and male snout vent length and condition (residuals of the regression between weight and snout vent length) was tested using a Generalised linear model with a Poisson error structure. Male display characteristics were also recorded. In *A. apletophallus* there are no courtship-specific behaviours, instead males have a repertoire of displays that are used in many contexts (Jenssen & Hover 1974). There are five different display types (A-F) and ‘A’ and ‘B’ displays are used most commonly (Jenssen & Hover 1974). The ‘A’ display involves full extension of the dewlap followed by a stereotypical head bob pattern (Jenssen & Hover 1974; Jenssen & Hover 1976). During the ‘B’ display the head bob sequence begins and the dewlap is extended on or after the first head bob is produced (Jenssen & Hover 1974). The type of display and the number of head bobs per display were recorded and compared between virgins and non-virgins using a Wilcox signed rank sum test. The ‘C’, ‘D’, ‘E’ and ‘F” displays are more commonly used in escalated contests between males and were not formally analysed in this study because they occurred at very low frequencies. All data were analysed using R.2.3.2 (R Development Core Team 2006). All individuals were released at the point of capture following experiments and the methods adhered to the ABS/ASAB guidelines for the ethical treatment of animals and were approved by the Smithsonian Tropical Research Institute (STRI) Institutional Animal Care and Use Committee.

## Results

### Experiment 1 – Male presented with a small virgin and a large non-virgin

Males displayed to the small virgin more often than to the large, non-virgin female (Figure 1, W = 108, *p* = 0.03). Males did not display more to the female that was closer at the beginning of the experiment (Binomial test *p* = 0.79). The total number of displays was not related to male snout vent length (F_1,11_ = 1.52, *p* = 0.24) or condition (F_1,11_ = 1.33, *p* = 0.27). The ‘A’ display was used most commonly, 43% of all displays were ‘A’ and 32.3% were ‘B’. The proportion of A to B displays did not differ when the male displayed to the virgin or the non-virgin (Wilcox signed rank sum test *p =* 0.59) and displays directed to a virgin did not differ in the number of head bobs compared to those directed to the non-virgin (mean number ± standard error (SE) of head bobs per display directed to virgin: 8.30±0.91, non-virgin: 10.67±2.71, Wilcox signed rank sum test *p =* 0.28).

**Figure 1.**
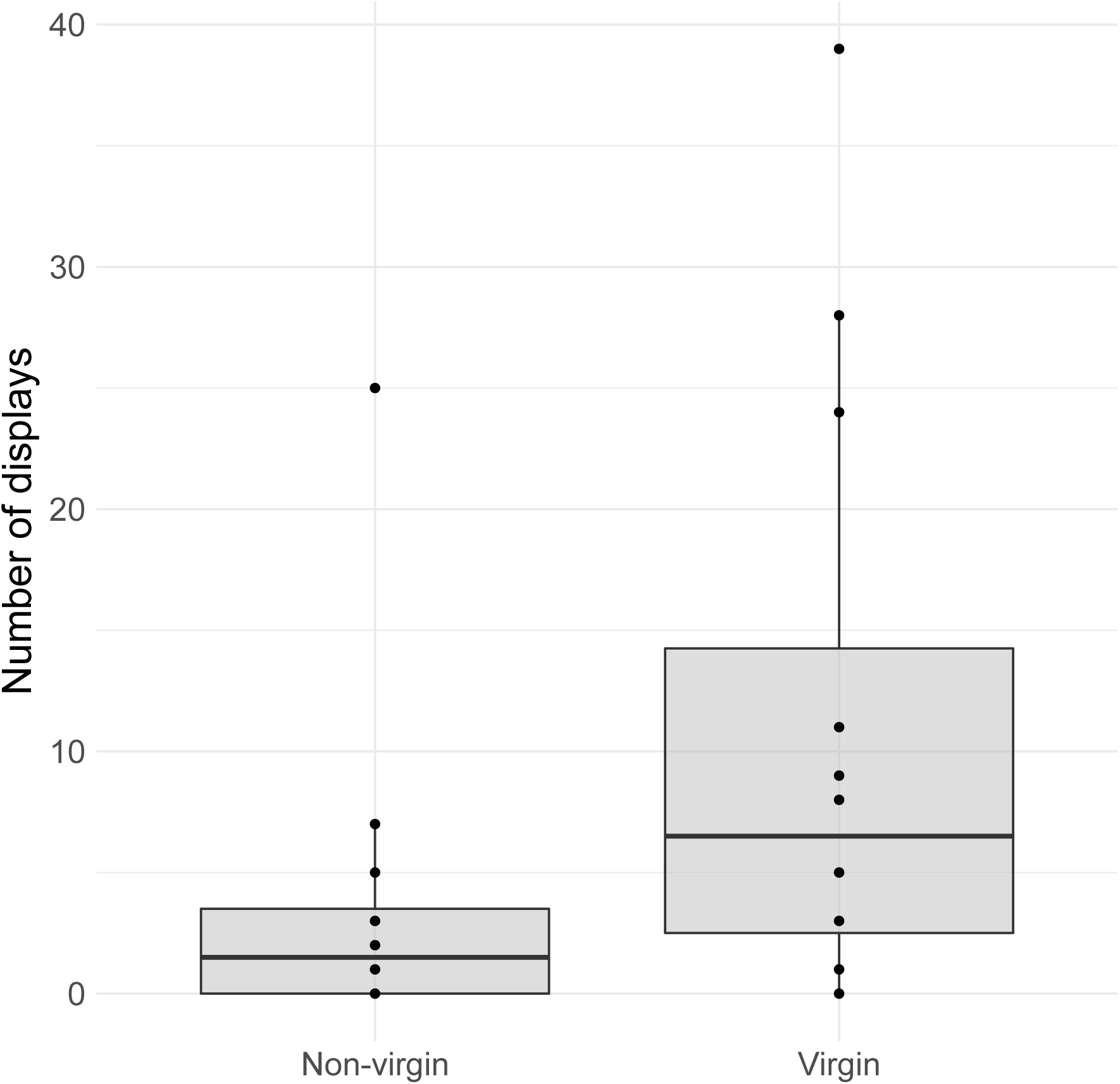
The mean number (± standard error) of male displays to the small virgin (40mm) and to the larger non-virgin female (45mm) during the 1 hour trial.

### Experiment 2 – Male presented with size matched females; one virgin, one non-virgin

Males displayed to virgins more often than non-virgins despite the fact that both females were the same size (Figure 2, W = 544.5, *p*<0.001). Similar to the first experiment, males did not display more to the female he was closest to at the beginning of the trial (Binomial test *p* = 0.77). The total number of displays was not related to male snout vent length (F_1,13_ = 0.13, *p* = 0.71) or condition (F_1,13_ = 0.47, *p* = 0.50). Again, males used ‘A’ displays most often (42.8%) and ‘B’ was the second most common display (33.9%), the proportion of ‘A’ to ‘B’ displays and the number of head bobs per display did not differ between displays directed to virgins and non-virgins (Proportion ‘A’ to virgin and non-virgin: Wilcox signed rank sum test *p* = 0.58; mean number ± SE of head bobs per display directed to virgin: 9.06±0.95, non-virgin: 7.51±1.09, Wilcox signed rank sum test *p =* 0.28).

**Figure 2.**
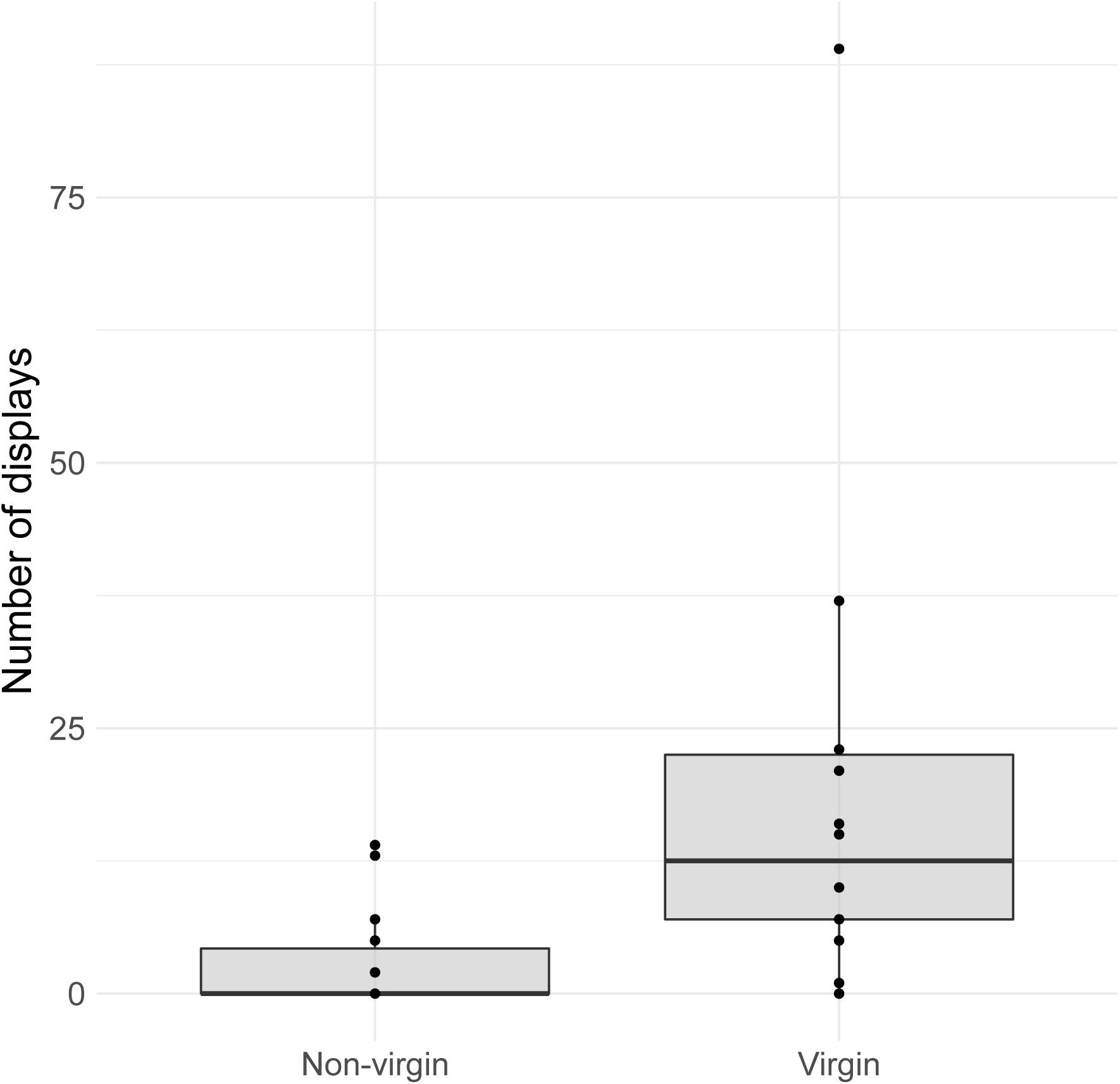
The mean number (± standard error) of male displays directed towards size matched virgin and non-virgin female during the 1 hour trial.

If I combine data from both experiments the findings do not change. The total number of displays in the second experiment was greater than the first (mean ± standard error number of displays: experiment 1 = 14.75 ± 4.75, experiment 2 = 22.9 ± 6.28), but this was not significant (Generalized linear model *F*_1,25_ = 1.06, *p* = 0.31).

## Discussion

When given a choice between a virgin and non-virgin female, males preferentially displayed to and courted the virgin female, even when the two females were the same size. This suggests that males preferentially court virgins and that they did not necessarily use body size as a cue of female mating history. In the field, body size would be an accurate cue to mating history as females >40mm SVL have sperm in their sperm storage tubules, whereas females <40mm SVL do not (Andrews 1989). In both experiments, there was no physical contact between the sexes so males could not use tactile cues and they could not sample chemical cues deposited on the female’s perches. Thus, it is likely they used visual cues to assess female mating history. In other cases where males prefer virgins, chemical signals were most commonly used (Bonduriansky 2001; Carazo et al. 2004), and in the only other vertebrate example, males were unable to determine female mating history using visual cues only (Guevara-Fiore et al. 2009). Thus, these results represent one of the few examples of male mate preference for virgin females based on visual cues.

This precopulatory male mating preference is likely to benefit males, by reducing sperm competition of the first few eggs laid by the female. The male that mates with the virgin is likely to have 100% paternity assurance of the first egg and depending on female mating frequency he may also sire several more eggs laid by the female in the absence of any sperm competition. As mentioned earlier, compared to other lizards, *A. apletophallus* seem reluctant to remate and have low level of multiple paternity (1-2 sires). In contrast in *Anolis sagrei*, 80% females mated multiply with up to four males siring offspring (Calsbeek et al. 2007), in *Eulamprus heatwolei*, multiple paternity in the field was 57% (Stapley & Keogh 2005) and in mating trials *Pseudemoia entrecasteuxii*, females were observed mating multiple times with the same male within a 10 hour period (Stapley 2008). Mating duration is long in *A. apletophallus* (mean copulation time = 94 minutes, range = 28-156 minutes) and if this is related to ejaculate volume or sperm transfer as it is in *Anolis sagrei* (Tokarz 1999) then it may represent a considerable energetic cost. In addition, courtship, involving dewlap extension and headbobs may increase vulnerability to predation and may be energetically demanding.

In contrast to expectations males did not use female body size to discriminate between virgins and non-virgins. In other cases where males prefer virgins, chemical signals were the most commonly used (Bonduriansky 2001; Carazo et al. 2004; Guevara-Fiore et al. 2009). In this study it is possible, but unlikely that males used chemical cues to assess female virginity. The female enclosures did no preclude airborne chemical cues, but males could not sample any substrate cues that females may have left on dowel perches. Anolis lizards, unlike some other lizard groups, rely almost completely on vision for communication and prey detection (Fleishman 1992). I never observed *A. apletophallus* tongue-flicking the air, but males were observed licking the dowel on a few occasions.

If males were not using chemical cues, how did they discriminate between virgins and non-virgins? In this system, I suggest that some aspect of female behaviour indicated her reproductive status. This could have involved changes in posture, head and dewlap movements and/or subtle changes in female dorsal colouration. Behavioural signalling of reproductive status has been demonstrated in other taxa, for example virgin females maybe less resistant to male advances (Guevara-Fiore et al. 2009) or may be more gregarious (Oku et al. 2005). Interestingly, male *A. apletophallus* appeared to make decisions about approaching and displaying to a female from a considerable distance (up to one metre), looking at both females and moving directly to the virgin and courting her. Therefore, it appeared that males used relatively long distance cues and he could discriminate between females before he began displaying. This suggests that it was not necessarily a difference in the response of the female to the male display that enabled him to discriminate. This is not to say that when the male approached and began to display intensely the female did not provided additional feedback the encouraged even further courtship. Unfortunately, it was not possible in this study to closely monitor the female’s behaviour and/or dorsal colouration but this may be worthy pursuit in future studies.

Male preference for virgin females was predicted to evolve under a certain set of circumstances, i.e. it was considered to be unlikely when female body size/age was positively related to reproductive success. However, results in the guppy contradict that and suggest this is not a general rule. More importantly, remating frequency and sperm storage may govern the evolution of this behaviour. In anoles and guppies females are able to store sperm between reproductive events, and they mate and lay eggs in a random order. In this sense, only in virgin females can males get any degree of paternity assurance and this assurance will decline with subsequent matings. Male preference for younger or smaller females could also represent a strategy to minimize sperm competition. Although relatively rare it has been observed in two other lizards groups (skinks (Stapley 2008) and chameleons (Stuart-Fox & Whiting 2005) and fish (Dosen & Montgomerie 2004), but in these cases avoidance of sperm competition was not the suggested explanation for the behaviour. It is plausible that because male preference for larger females is so pervasive in vertebrates, male preference for virgins is relatively rare. The results of this study, highlight that in species where large females do not have reproductive benefits, or when sperm is stored between subsequent reproductive events selection on male preference for virgins may exist and give rise to the evolution of this preference function.

## Acknowledgements

Special thanks go to Raineldo Urriola and Argeli Ruiz for logistic support in Panama. Thanks also go to John Christy for advice and support. Funding was provided by the Smithsonian Tropical Research Institute.

